# Insights on long-distance dispersal, ecological and morphological evolution in the fern genus *Microgramma* from phylogenetic inferences

**DOI:** 10.1101/2020.06.07.138776

**Authors:** Thaís Elias Almeida, Alexandre Salino, Jean-Yves Dubuisson, Sabine Hennequin

**Affiliations:** Herbário HSTM, Universidade Federal do Oeste do Pará, Av. Marechal Rondon, s.n. – Santarém, Pará, Brazil. CEP 68.040-070; Departamento de Botânica, Universidade Federal de Minas Gerais, Av. Antônio Carlos, 6627 – Belo Horizonte, Minas Gerais, Brazil. Caixa Postal 486, CEP 30123-970; Institut de Systématique, Evolution, Biodiversité (ISYEB), Sorbonne Université, Muséum national d’Histoire naturelle, CNRS, EPHE. Université des Antilles, 57 rue Cuvier, 75005 Paris, France

**Keywords:** ant-fern association, biogeography, frond dimorphy, ferns, homoplasy, long-distance dispersal

## Abstract

The epiphytic fern genus *Microgramma* (Polypodiaceae) comprises 30 species occurring mainly in the Neotropics with one species in Africa, being an example of trans-Atlantic disjunction. Morphologically and ecologically, *Microgramma* presents a wide variation that is not seen in its closest related genera. Recent works changed the circumscription of *Microgramma* to better conform with phylogenetic evidence, but no comprehensively sampled study has addressed the evolution of this lineage. This study aimed to investigate phylogenetic relationships, ecological and morphological evolution within *Microgramma*, as well as test the role of long-distant dispersal in the history of the genus. Sequences from five plastid regions were used to infer the phylogenetic relationships and estimate divergence times. Our results show five clades in *Microgramma* that do not corroborate any infrageneric classification system proposed. Several morphological traits seem to be homoplastic, such as leaf dimorphism. Tuber-like myrmecodomatia are suggested to be synapomorphic for one clade, although ant-plant association appears in two distinct lineages. *Microgramma lycopodioides* and *M. mauritiana* are not closely related, with the African species nested within an Atlantic Forest clade, indicating a long-distance dispersal event estimated to have occurred around 15 Ma from South America to Africa, followed by speciation.

## INTRODUCTION

*Microgramma* C.Presl is a fern genus of approximately 30 species in the family Polypodiaceae, all occurring in the Neotropics except one occurring in Africa. They are rhizome-creeping epiphytes, growing from trunk bases to the canopy, with a few species also able to grow in rocky or terrestrial habitats. They occur preferentially in wet tropical forests but also colonize dry forests and savanna environments across a broad area of South America and tropical Africa (Almeida, 2020). *Microgramma* is part of the neotropical clade of Polypodiaceae and it is placed in the campyloneuroid clade with *Campyloneurum* C.Presl and *Niphidium* J.Sm. (Schneider *et al*., 2004; Labiak & Moran, 2018). Although its relationships with other genera are well known, the phylogenetic tree of *Microgramma* itself has not been previously explored.

*Microgramma* presents a broader spectrum of frond and rhizome variation than is found in its closest related genera (Schneider *et al*., 2004; Salino *et al*., 2008; Labiak & Moran, 2018). Two different functional fronds are found in *Microgramma*: fertile fronds bearing sporangia, and sterile fronds with the primary purpose of photosynthesis (Wagner & Wagner, 1977). Monomorphic species bear fertile and sterile fronds that are similar in size and shape, while dimorphic species have fertile fronds that differ from sterile fronds in size, or size and shape. The rhizomes range from cylindric to flattened, and can have tubers, sac-like myrmecodomatia that are described to have associations with ants (Gómez, 1974; Salino *et al*., 2008).

Regarding geographic distribution, *Microgramma*, along with *Pleopeltis* Humb. & Bonpl. ex Willd., is unique in the neotropical Polypodiaceae clade (outside the grammitids) in being an example of disjunction between Africa and South America. Its species occur throughout the Neotropics, from Florida to Argentina, with a single species [*Microgramma mauritiana* (Desv.) Tardieu] in Sub-Saharan Africa, the Malagasy region, and the Indian Ocean Islands (Moran & Riba, 1995; Roux, 2009; Smith *et al*., 2018). Given their morphological similarity, populations from both continents were once considered conspecific under a broadly distributed *M. lycopodioides* Copel.; but African-Malagasy populations of *Microgramma* are now recognized as *M. mauritiana*, while *M. lycopodioides* is restricted to the Neotropics (Salino *et al*., 2008; Roux, 2009). In the Neotropics, some species are widespread *[M. percussa* (Cav.) de la Sota and *M. lycopodioides* (Smith *et al*., 2018)], with relatively few narrow-range endemics such as *M. rosmarinifolia* (Kunth) R.M.Tryon & A.F.Tryon (Tryon & Stolze, 1993), *M. recreense* (Hieron.) Lellinger (León & Jørgensen, 1999), and *M. crispata* (Fée) R.M.Tryon & A.F.Tryon (Almeida, 2020).

Long-distance dispersal (LDD) is a key process in shaping plants’ current distribution in natural environments (Jordano, 2017). Many examples from different plant lineages document the importance of LDD in shaping floristic relationships between Africa and South America (see Christenhusz and Chase 2013 for a review). Vascular seedless plants are lineages where it is expected to find very closely related populations or species occurring disjunctly (Tryon, 1986). Their life cycle have two alternating generations - sporophyte and gametophyte - which are heteromorphic and independent (Haufler *et al*., 2016; Pinson *et al*., 2017), and their small spores are easily dispersed by the wind (Sheffield, 2008). Moran & Smith (2001) documented several examples of LDD in ferns and lycophytes between Africa and South America based on the morphological similarity of species (hypothesized as species pairs) or populations occurring in both continents (hypothesized to be conspecific). Access to phylogenetic data now allows us to test these hypotheses. A few of them were not corroborated in previous studies (e.g. Rouhan *et al*., 2004; Lehtonen *et al*., 2010; Link-Pérez, Watson, & Hickey, 2011; Labiak *et al*., 2014; Schuettpelz *et al*., 2016; Huiet *et al*., 2018) while many were found to be the result of LDD (e.g. (Prado *et al*., 2013; Almeida *et al*., 2016; Gasper *et al*., 2016; Bauret *et al*., 2017, 2018; Duan *et al*., 2017).

Even though recent studies changed the circumscription of *Microgramma* to better conform phylogenetic evidence (Salino *et al*., 2008; Almeida *et al*., 2017), no comprehensively sampled study has, to the date, addressed the morphological variation, the ant-fern-association, and the disjunction between neotropical and African populations within the genus. This study aimed to investigate phylogenetic relationships, ecological and morphological evolution within the genus *Microgramma* as well as test the role of long-distant dispersal in the history of the genus.

## MATERIAL AND METHODS

### Taxon sampling

Twenty-six out of the ca. 30 species of *Microgramma* were sampled (105 newly acquired sequences), covering all known putative species groups as well as the geographic range of the genus. Species not sampled were mostly known from single locations (*Microgramma recreense*), are poorly known or rare taxa (*M. ulei* (Ule) Stolze and *M. tuberosa* (Maxon) Lellinger) or are known only for their type specimen (*M. fosteri* B.León & H.Beltrán). Two exemplars per species were included and within species, samples were selected to encompass the geographic or morphological variation of the known species.

Outgroups were selected from eight Polypodiaceae genera based on the studies of Schneider *et al*. (2004) and Almeida et al. (2017) and include nine species of *Campyloneurum*, three of *Niphidium* and one species of *Adetogramma* T.E.Almeida, *Pecluma* M.G.Price, *Phlebodium* (R.Br.) J.Sm., *Pleopeltis, Polypodium* L., and *Serpocaulon* A.R.Sm. (Appendix). The samples were stored in silica gel and vouchers were incorporated to the following herbaria: BHCB, FURB, LPB, PMA, USM, USZ (acronyms follow Thiers 2020 onward: http://sweetgum.nybg.org/science/ih/). Vouchers and GenBank accessions are listed in the Appendix. Aligned data matrix was deposited in TreeBASE (http://purl.org/phylo/treebase/phylows/study/TB2:S26342).

### Sequences acquisition

Total DNA was extracted from field-acquired silica gel-dried or fresh tissues, using the Qiagen DNeasy Plant mini kit (Qiagen Inc., Valencia, CA, USA). PCR amplifications were performed for five plastid regions: *rbcL, rps4*, (*rps4* gene, and *rps4-trnS* intergenic spacer), and *trnLF, (trnL* intron, *trnL-trnF* IGS and including short exon portions of the genes *trnL* and *trnF*). Amplifications were done in single reactions with primers 1F and 1365R (Haufler & Ranker, 1995) for the *rbcL* region, the primers rps5F (Nadot *et al*., 1995) and trnSR (Smith & Cranfill, 2002) for the *rps4* gene and *rps4-trnS* IGS, and the primers Fern1 (Trewick *et al*., 2002) and f (Taberlet *et al*., 1991) for the *trnL* intron, and *trnL-trnF* IGS. Polymerase chain reactions were performed in a 20 μL solution containing 1.0 μL of genomic DNA template, 2.0 μL of PCR buffer (Qiagen 10x PCR Buffer), 1.0 μL of DMSO, 1.0 μL of BSA (4 mg/mL), 0.8 μL of dNTPS (10 mM), 0.32 μL (10 μM) of each primer, 0.12 units of Taq DNA polymerase (Qiagen, 5 units/μL) and 14.44 μL of ultra-pure water. The thermal cycling conditions were the same for all three regions: 3 min at 94 °C, 35 cycles of 45 s at 94°C, 60 s at 53°C and 90s at 72°C, followed by 5 min at 72 °C. The amplicons were purified by precipitation with polyethylene glycol - PEG and sequenced by Macrogen (Seoul, South Korea) for bidirectional sequencing reaction in an ABI3730XL.

### Alignment and phylogenetic analyses

Sequence electropherograms were edited using the Geneious version R11 (http://www.geneious.com, Kearse et al. 2012). The edited sequences were submitted to automated alignment with MUSCLE and the resulting alignment was checked manually using MEGA 7 (Kumar, Stecher, & Tamura, 2016).

The data were analyzed using maximum likelihood (ML) and Bayesian inference (BI). We inferred the Maximum likelihood tree using the IQ-TREE webserver (Nguyen *et al*., 2015; Trifinopoulos *et al*., 2016) with a partitioned matrix, automatic selection of the best-fit substitution model using Bayesian Information Criterion (Schwarz, 1978) through ModelFinder (Kalyaanamoorthy *et al*., 2017), and with branch support assessed using the ultrafast bootstrap approximation with 1,000 bootstrap replicates (Minh, Nguyen, & Von Haeseler, 2013; Hoang *et al*., 2018). The partitions were selected by Partition Model (Chernomor, von Haeseler, & Minh, 2016). For BI we performed a model-based Markov chain Montecarlo-based phylogenetic analysis using MrBayes v3.2.2 (Ronquist *et al*., 2012) through Cipres Science Gateway (Miller, Pfeiffer, & Schwartz, 2010), treating each DNA region (*rbcL, rsp4* gene, *rps4* IGS, and *trnL-trnF*) as separate partitions. An evolutionary model for each DNA region was selected in jModelTest 2 (Guindon & Gascuel, 2003; Darriba *et al*., 2015), using the Bayesian Information Criterion (Schwarz 1978, Table 1). Each analysis consisted of two independent runs with four chains performing 50,000,000 generations, sampling one tree every 1,000 generations. For the BI and ML, GenBank sequences of *Polypodium vulgare* L. were used as the outgroup. After discarding the first 10% of the trees as burn-in, the remaining trees were used to assess topology and posterior probabilities (PP) in a majority-rule consensus. To check the convergence of the runs, ESS (effective sample size) and PSRF (potential scale reduction factor) were examined (Ronquist *et al*., 2012) using Tracer v.1.6 (Bouckaert *et al*., 2014). Based on the sampled parameter values examined, 10% of the trees, including the ones generated during the burn-in phase, were discarded. Remaining trees were used to assess topology and posterior probabilities (PP) in a majority-rule consensus tree. Results were summarized on a majority rule consensus tree.

**Table 1:**
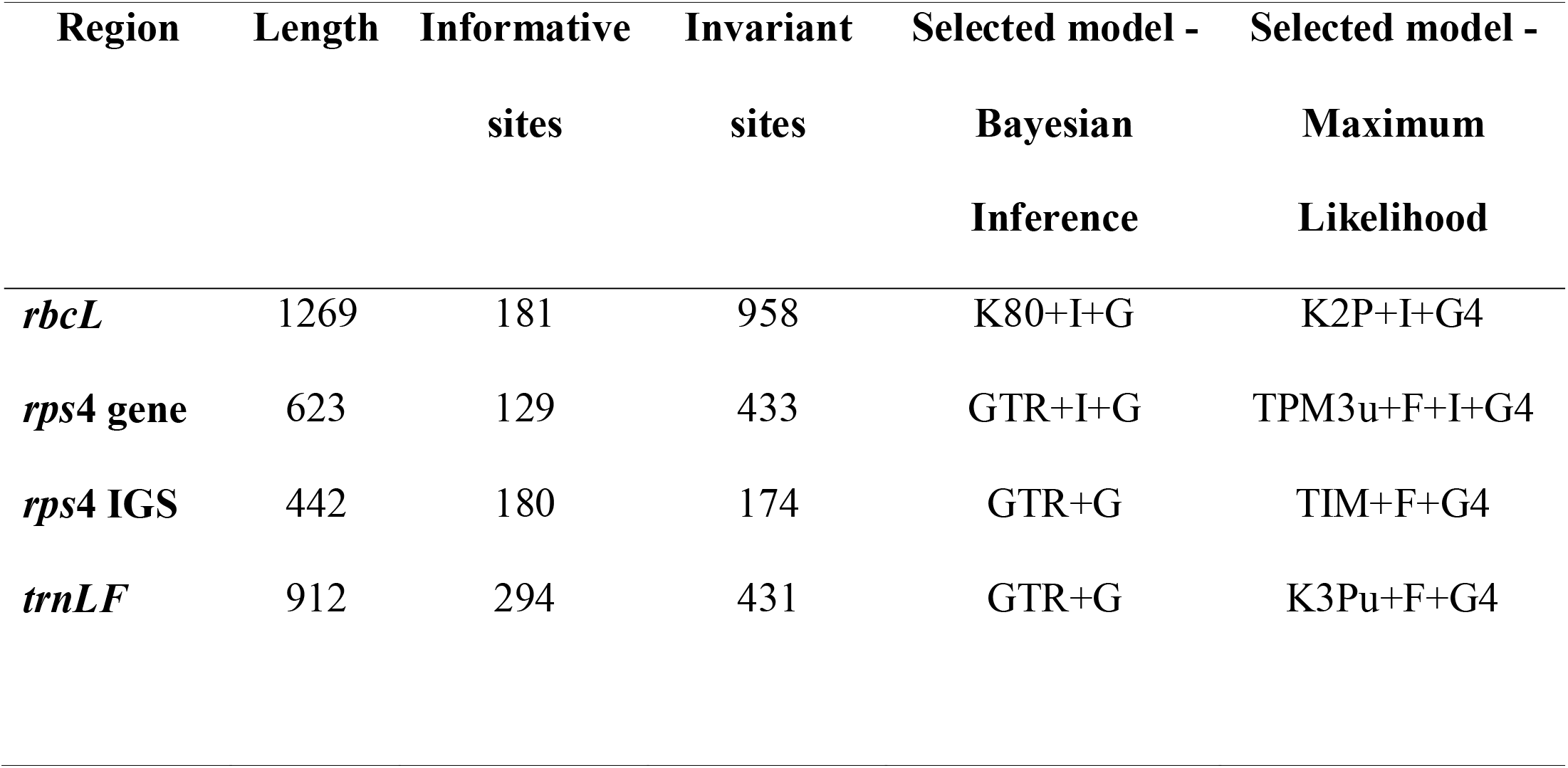
Size, number of informative sites and nucleotide substitution models selected for each partition used in this study

### Divergence times estimations

Divergence times were estimated with the software BEAST 2.5 (Bouckaert *et al*., 2019) through the CIPRES Science Gateway (Miller *et al*., 2010). The data matrix was partitioned among the four markers and assigned the GTR model of substitution for the *rps4* gene, *rps4* IGS, and *trnL-trnF* intron+IGS, and the HKI model for the *rbcL* gene. The trees and clock models were linked. In the absence of reliable fossil for the ingroup, we used time estimations from the large-scale study on the time diversification of leptosporangiate ferns by Testo and Sundue (2016) for node calibration. We used calibrations for the following nodes with the respective ages: campyloneuroid clade + *Adetogramma* + *Serpocaulon* = 55.84 ma; campyloneuroid clade = 49.54 ma; *Campyloneurum* + *Niphidium* = 45.64 ma; *Campyloneurum* = 35.28 ma; *Microgramma* = 26.73 ma; *Niphidium* = 5.97 ma. We assigned a normal distribution for all calibrated nodes. We used an uncorrelated lognormal clock, and the birth-death tree prior, with the relative extinction rate set to 0.5 and speciation set for 1.0. Parameters and priors were set using BEAUti (Drummond *et al*., 2012). A starting tree topology was specified based on the resulting majority-rule Bayesian tree, using the package ape (Paradis & Schliep, 2019) in R (R Core Team, 2019) to match branch lengths to units of time specified for BEAST analysis.

Two Markov chain Montecarlo runs were conducted using 5 × 10^7^ generations with parameters sampled every 5,000 steps. Results from both BEAST runs were analyzed to confirm that stationarity, convergence, and effective sample sizes (ESS) were sufficient for all parameters using Tracer 1.7 (Rambaut *et al*., 2018). Trees were summarized and annotated with median ages estimates and 95% highest posterior density (HPD) intervals using LogCombiner and TreeAnnotator in BEAST 2 (Bouckaert *et al*., 2019), with 20% of trees discarded as burn-in. A tree was generated from the remaining 8,000 trees using the BI majority-rule tree as a target tree to constrain the topology and for the dated tree to reflect the polytomies found in both BI and ML analyses.

## RESULTS

The length of each sampled region, number of parsimoniously informative and invariant sites, and models used for Bayesian Inference and Maximum Likelihood analyses are given in Table 1. In both the Bayesian Inference and BEAST analyses, the two independent runs converged and ESS values for all parameters were higher than 200, indicating convergence of the analysis (Barido-Sottani *et al*., 2018).

### Phylogenetic analyses

The monophyly of *Microgramma* was strongly supported (1.00 PP, 100% BS) (Figure 1). Four main clades are recognized inside *Microgramma* (Figure 1). The Andean endemic *Microgramma rosmarinifolia* is recovered as sister to the remaining species of *Microgramma* (1.00 PP, 100% BS). At the subsequent node, a clade called here the Scaly clade, is sister to all remaining clades (0.98 PP, 97% BS). The Scaly clade includes all species with subulate scales (*M. latevagans* (Maxon & C.Chr.) Lellinger, *M. nana* (Liebm.) T.E.Almeida, *M. piloselloides* (L.) Copel.,*M. reptans* (Cav.) A.R.Sm.,*M. tecta* (Kaulf.) Alston,*M. tobagensis* (C.Chr.) C.D. Adams & Baksh.-Com.) and round scales (*M. dictyophylla* (Kunze ex Mett.) de la Sota and *M. percussa*) covering both sides of fronds. These species are distributed throughout the Neotropics, with widespread species such as *M. nana*, *M. percussa*, *M. reptans*, and *M. tobagensis*; species with restricted distribution as *M. dictyophylla* (Northern South America), and *M. piloselloides* (Central America and Caribbean), as well as endemics, such as *M. latevagans* (Bolivia and Peru), and *M. tecta* (Atlantic Forest). In the following divergence, the predominantly Eastern South American *M. squamulosa* (Kaulf.) de la Sota (including the supposedly hybrid species *M. mortoniana* de la Sota) appears as sister to all the remaining species (1.00 PP, 100% BS). Subsequently, there is a polytomy including three clades: the Vacciniifolia clade, Lycopodioides clade, and Persicariifolia clade. The Vacciniifolia clade (0.79 PP, 59% BS) includes *M. mauritiana, M. crispata, M. geminata* (Schrad.) R.M.Tryon & A.F.Tryon, and *M. vacciniifolia* (Langsd. & Fisch.) Copel. The African *Microgramma mauritiana* appears nested in this lineage, although the position is not supported by the ML analyses. The remaining species in this clade are Atlantic Forest endemics (*M. crispata* and *M. geminata*) and the widespread *M. vacciniifolia* which occurs in the Atlantic Forest and also has disjunct populations in northern South America, the Caribbean, eastern Andes, and the Chaco.

**Figure 1.**
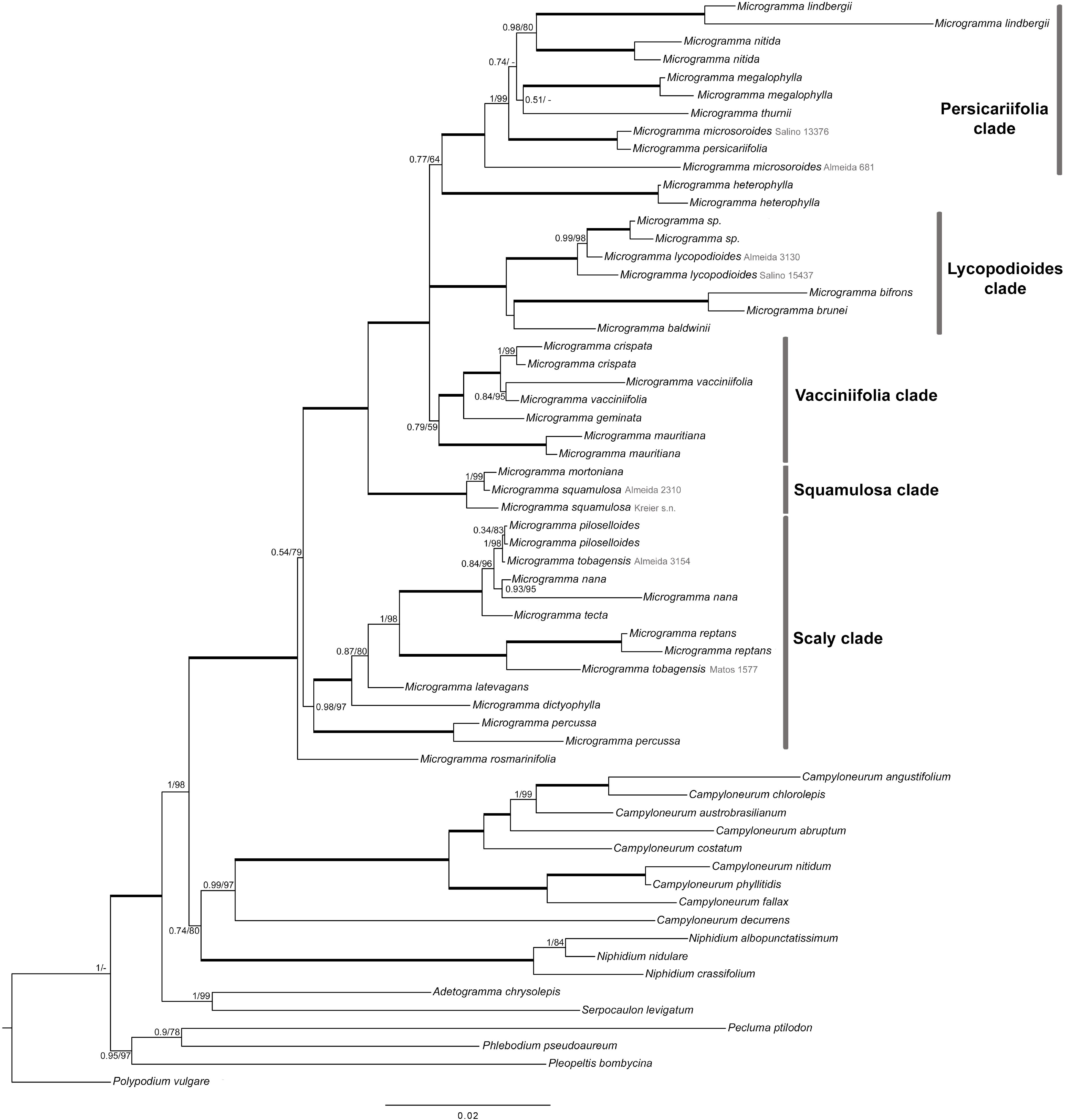
Majority rule consensus tree resulting from Bayesian inference analyses from the *rbcL, rps4*, and *trnLF* datasets. Values next to branches nodes correspond to Bayesian posterior probabilities and maximum likelihood bootstrap support, respectively. Thickened branches indicate those supported with 1.00 PP and 100% BS. Clades names are as indicated in the text. Vouchers are indicated for those species that are not monophyletic.

The Lycopodioides clade (1.00 PP, 100% BS) includes the widespread neotropical *M. lycopodioides* in a polytomy with *M. baldwinii* Brade and a clade including the representatives of myrmecophytic species (*M. bifrons* (Hook.) Lellinger and *M. brunei* (Wercklé ex Christ) Lellinger) (1.00 PP, 100% BS). We found no support for the recognition of African populations, recognized here as *M. mauritiana*, to be related to or conspecific with *M. lycopodioides*. The morphological similarities between these species are likely to be homoplastic.

The Persicariifolia clade (0.77 PP, 64% BS) includes the Mesoamerican *M. nitida* (J.Sm.) A.R.Sm., the eastern South American *M. lindbergii* (Mett. ex Kuhn) de la Sota and *M. microsoroides* Salino, T.E.Almeida & A.R.Sm., the lowland Amazonian *M. megalophylla* (Desv.) de la Sota and *M. thurnii* (Baker) R.M.Tryon & Stolze, and the widespread *M. persicariifolia* (Schrad.) C.Presl. The Central American and Caribbean species *M. heterophylla* (L.) Wherry was sister to other species in the Persicariifolia clade, with its position poorly supported in both BI and ML analyses (0.77 PP, 64% BS) and therefore is not considered within this clade.

The ant-associated species *M. bifrons* and *M. brunei*, representatives of the group previously segregated as the genus *Solanopteris* Copel. (along with the unsampled *M. tuberosa* and *M. fosteri*) do not belong to the same clade as the only other known ant-associated species *M. megalophylla* (Figure 1). Additionally, leaf monomorphism is recovered as symplesiomorphic and leaf dimorphism occurs in a derived position in species distributed in all four clades.

The inclusion of more than one terminal for most species indicate that some might not be monophyletic: *M. microsoroides* is recovered as paraphyletic to *M. persicariifolia*, *M. mortoniana* is nested in *M. squamulosa*, and *M. tobagensis* is recovered as polyphyletic (Figure 1).

### Divergence times analysis

*Microgramma rosmarinifolia* was estimated to diverge around 26.85 Ma (95% HPD 25.89– 27.84 Ma), succeeded by the divergence of the Scaly clade around 26.48 Ma (95% HPD 24.11–27.76 Ma) (Figure 2). The divergence of *M. squamulosa* was estimated to have occurred at 22.89 Ma (95% HPD 19.5 – 25.92 Ma). We do not present the dates from the polytomy including *M. heterophylla*, the Vacciniifolia, Lycopodioides, and Persicariifolia clade, to conform with the results from the BI and ML. The divergence of *M. heterophylla* from the Persicariifolia clade was estimated at 19.73 Ma (95% HPD 13.62 – 22.1 Ma). The divergence of the Vacciniifolia clade was estimated to have occurred at 15.07 Ma (95% HPD 9.29–19.69 Ma), and that of the Lycopodioides clade and the Persicariifolia clade was estimated to have occurred at 12.98 Ma (95% HPD 8.15–17.68 Ma) and 15.22 Ma (95% HPD 10.85–19.63 Ma), respectively.

**Figure 2.**
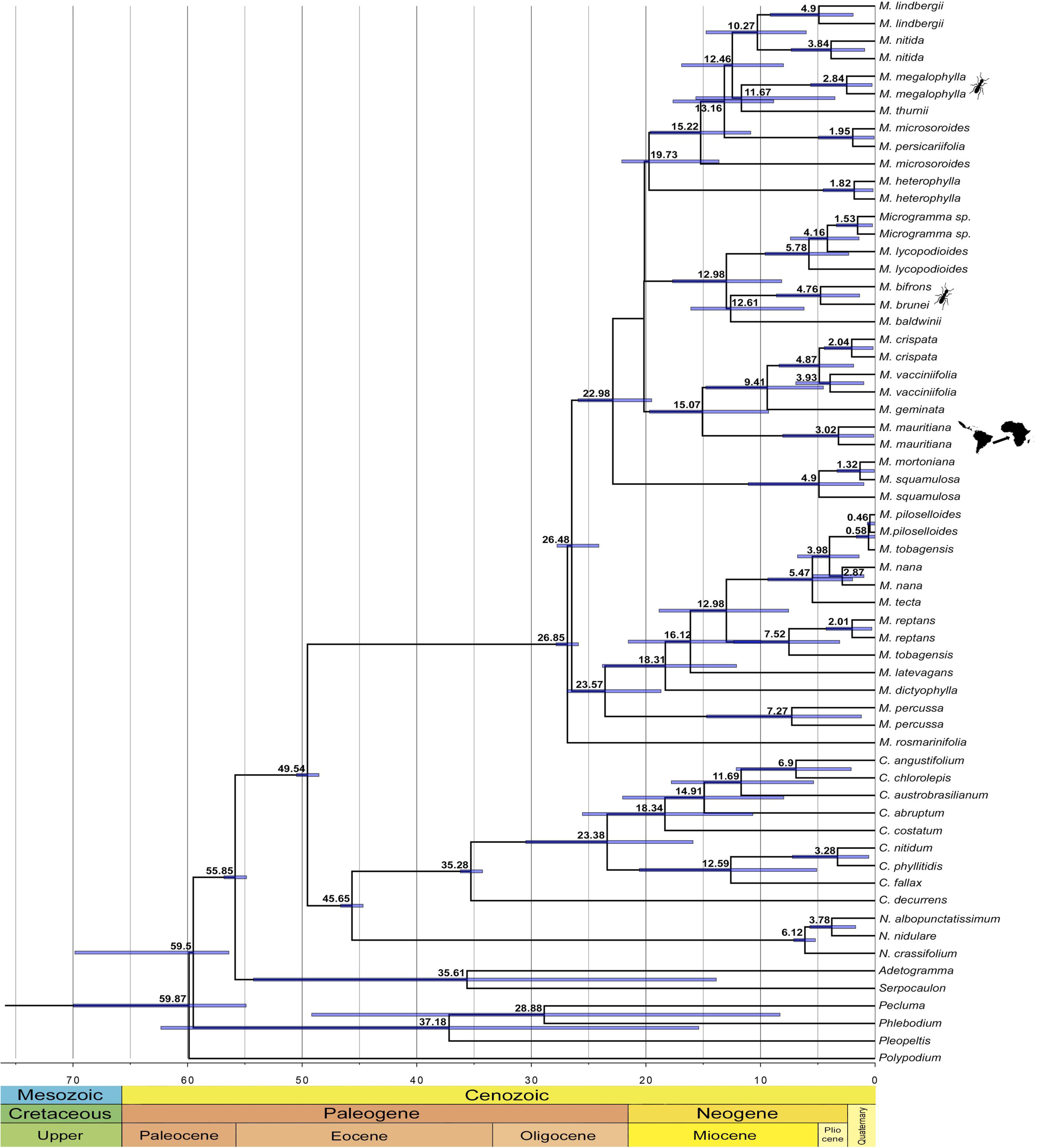
Divergence time estimates for *Microgramma* inferred from the *rbcL, rps4*, and *trnLF* datasets. The ant figures indicate the lineages displaying records of ant-fern. The map outline indicates the lineage derived from a long-distance dispersal event from South America to Africa. Blue lines at the nodes indicate 95% high probability density intervals. Dates are given in million years (Ma).

The lineage containing the species with myrmecodomatia (*M. brunei*, *M. bifrons*) (Figure 3), was estimated to have diverged from *M. baldwinii* (species with no records of ant-fern association) around 12.61 Ma (95% HPD 6.2 –16.08 Ma). *Microgramma megalophylla*, another species with ant-fern association that does not have myrmecodomatia (Figure 3), was estimated to have diverged from *M. thurnii* (species with no records of ant-fern association) ca. 11.67 Ma (95% HPD 3.5–15.64 Ma) (Figures 1, 2).

**Figure 3.**
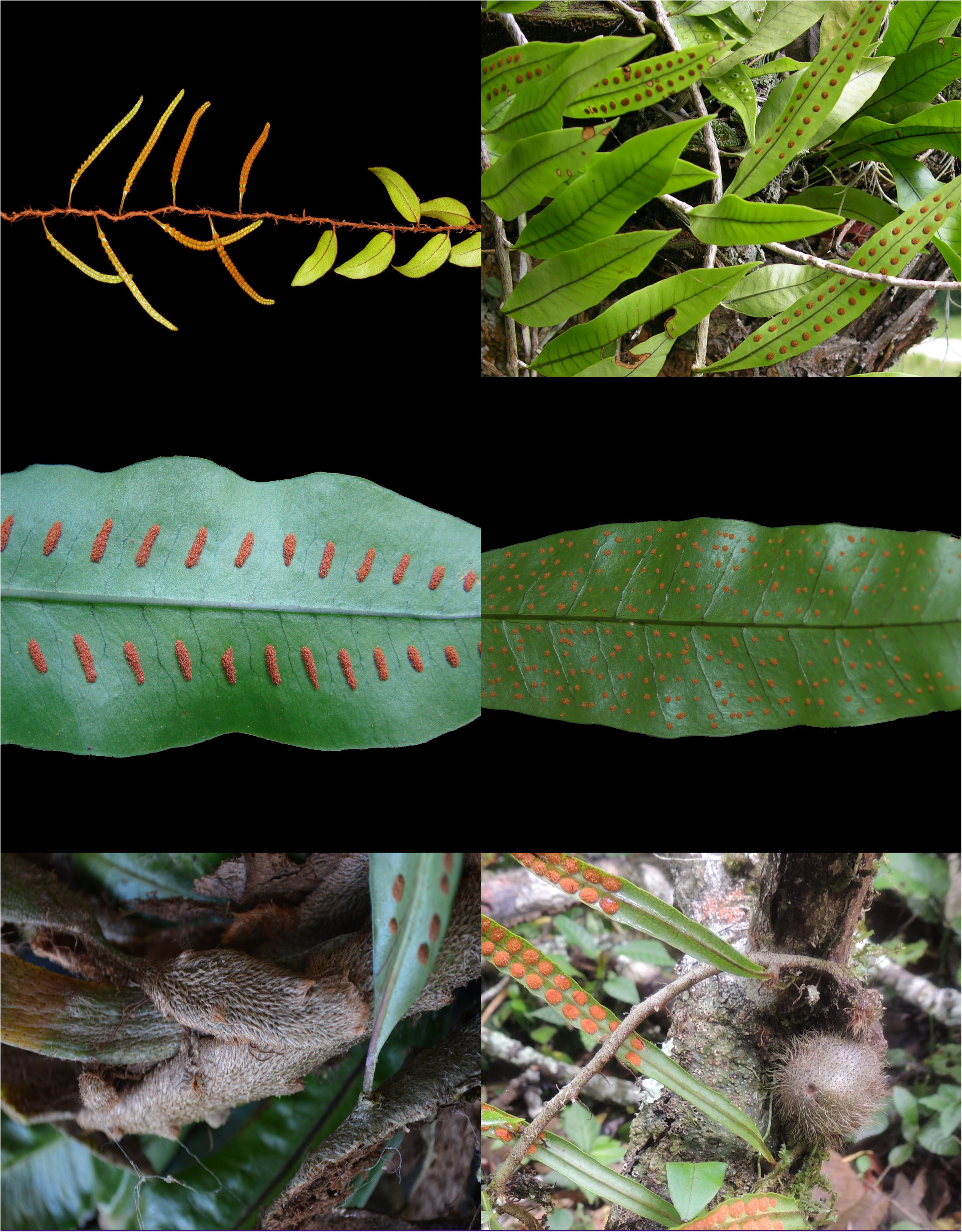
Images illustrating characters discussed in the text. A – Frond dimorphism in *M. reptans*. B – Frond monomorphism in *M. geminata*. C. Linear sori in *M. persicariifolia*. D – Scattered irregular sori in *M. microsoroides*. E – Flattened rhizomes in *M. megalophylla*. F – Cylindrical rhizomes with tubers in *M. brunei* (Photos: A – C.N. Fraga, C – N.F.O. Mota, D, F – A. Salino).

## DISCUSSION

### Phylogenetic relationships within *Microgramma*

Our results support the circumscription of *Microgramma* proposed by Salino et al. (2008), (based on approximately one third of the genus’ species sampled) including the species previously segregated in *Solanopteris* [*M. bifrons, M. brunei, M. fosteri*, and *M. tuberosa* (=*M. bismarckii* (Rauh) B.Léon)], and also *M. percussa* and *M. dictyophylla* (Figure 1). These results do not support generic classifications that split the genus and do not corroborate any infrageneric classification previously proposed for the genus. For example, the recognition of *Anapeltis* J.Sm. and *Craspedaria* Link proposed by Christensen (1938), Ching (1940), and Pichi Sermolli (1977). The characters used to segregate *Craspedaria* were venation type, presence or absence of included free veinlets directed towards the costa, shape of sori, and the presence or absence and shape of paraphyses (de la Sota, 1973a; Pichi Sermolli, 1977). These features are variable among *Microgramma* species and may constitute homoplastic characters, thus possibly devoid of any systematic value for use at the generic level. de la Sota (1973) suggests, for example, that the evolutionary process of venation simplification may be related to the reduction of leaf size and increase of its dimorphism. Our data indicates these events have occurred independently several times among species of the Scaly clade (such as *M. reptans*), Vacciniifolia clade (*M. vacciniifolia*), Lycopodioides clade (*M. bifrons*), and *M. squamulosa* (Figure 1).

The recognition of *Solanopteris* was mainly based upon the presence of tubers, although Copeland (1951) cites fronds texture and vein and veinlet pattern as characters that could be used to recognize this group of species in a separate genus. Conversely, Lellinger (1977), when combining *Microgramma* subg. *Solanopteris*, stated that the features used by Copeland (1951) were insufficient to grant the group a generic status. The presence of echinate spores in the myrmecophytic species was another feature used to segregate *Solanopteris* from *Microgramma* (Tryon & Lugardon, 1991). Those characters (tubers and echinate spores) could be synapomorphies of the clade. The inclusion of the other myrmecophytic species with tubers, *M. fosteri* and *M. tuberosa*, is needed to test this hypothesis.

In addition, our results do not corroborate the infrageneric classification system proposed by Lellinger (1977). The presence of scales on the fronds, a character used by Lellinger (1977) to group species in *Microgramma* subg. *Lopholepis* (J.Sm.) J.Sm. (including *M. piloselloides, M. reptans*, and *M. vacciniifolia*), is a homoplastic feature. Scales are present in species that appear in several clades: *M. piloselloides* and *M. reptans* in the Scaly clade and *M. vacciniifolia* in the Vacciniifolia clade (Figure 1). Moreover, the scales present in most species from the Scaly clade greatly differ in shape from the scales of *M. vacciniifolia* and *M. squamulosa* and are probably homoplastic.

### Morphological characters

The presence of myrmecodomatia in *Microgramma* species that are not inferred as closely related (Figure 1) indicates either two origins, or one origin and at least five losses. Within *Microgramma*, two forms of myrmecodomatia have been reported: the broad domed rhizomes enclosing a cavity below recently found in *Microgramma megalophylla* (Almeida, 2018) and the more specialized chambered tubers well known for *Microgramma bifrons* and *M. brunei* (Gómez, 1974). The unique rhizome form of *M. megalophylla* resembles a simple form of what is found in the paleotropical ant-fern genus *Lecanopteris* (Gay, 1993; Almeida, 2018) (Figure 3). *Lecanopteris mirabilis* (C.Chr.) Copel. has the most basic form of myrmecodomatia present in that genus and is sister to other *Lecanopteris* that have more complex chambered myrmecodomatia (Gay, 1993; Haufler *et al*., 2003; Testo *et al*., 2019). In addition to myrmecodomatia, the spores of *M. bifrons* and *M. brunei* are echinate, a feature proposed to be associated with ant dispersal (Hennipman, 1990; Tryon & Lugardon, 1991), but this feature is absent from *M. megalophylla* (Tryon & Lugardon, 1991). These two forms of ant-association belong to two different clades, evidence supporting a hypothesis of two separate origins. This parallel between *Lecanopteris* and ant-associated *Microgramma* is consistent with the hypothesis proposed by Almeida (2018) that flattened domed rhizomes are the most basic form in a transition to more complex chambered domatia (Haufler *et al*., 2003; Testo *et al*., 2019).

Frond dimorphism in *Microgramma* appears to be homoplastic (Wagner & Wagner, 1977), occurring in all clades: *M. nana*, *M. reptans*, *M. tecta*, and *M. tobagensis* in the Scaly clade; *M. crispata*, *M. mauritiana*, and *M. vacciniifolia*, in the Vacciniifolia clade; *M. bifrons*, and *M. brunei* in the Lycopodioides clade, and *M. nitida* in the Persicariifolia clade. This feature is unique among the Campyloneuroid clade (formed by *Microgramma+Campyloneurum+Niphidium* - Schneider *et al*., 2004), and rare among the neotropical-clade of Polypodiaceae, being found mostly in the genus *Pleopeltis* (Smith & Tejero-Díez, 2014). The derived occurrences of frond-dimorphism within *Microgramma* is evidence that frond monomorphism is symplesiomorphic in the Campyloneuroid clade. Although they can be advantageous during reproduction (e.g. more production and effective dispersal of spores) (Wagner & Wagner, 1977), dimorphy can have an impact on plants because the reduced fertile fronds are generally short-lived (Mehltreter & Sharpe, 2013) and have lower photosynthetic rates, suggesting an elevated cost to be produced and maintained (Wagner & Wagner, 1977; Watkins, Churchill, & Michele Holbrook, 2016). Further studies are needed to clarify if, from a developmental point of view, frond dimorphism is homologous throughout *Microgramma*, and which selective pressures and modifications in developmental processes could have induced or facilitated so many transitions from monomorphy to dimorphy.

Despite the above-discussed scenario of homoplasies, some distinctive morphological characters traditionally used to diagnose species and species relationships in *Microgramma* may be synapomorphies. For instance, the Scaly clade comprises species with subulate scales in the fronds and also includes *M. dictyophylla*, a species that presents minute, round, dot-like scales covering the fronds surface, and *M. percussa*, with round scales spread across both surfaces of the fronds (Almeida, 2014). The distribution pattern of the scales in the fronds of *M. percussa* and *M. dictyophylla* are similar to the species with subulate scales. Also, in the Persicariifolia clade, which aggregates the largest species in the genus, species are usually found as epiphytes on tree trunks but keep contact with soil through rhizome or pending roots (hemiepiphytes), (Testo & Sundue, 2014), e.g. *Microgramma persicariifolia*. However, given the morphological variation found in *Microgramma*, further analyses are needed to properly define 1) the homology among features and to 2) understand the evolution of morphological characters in the genus.

### Possible issues with species circumscriptions

Although the circumscription of species was not the focus of this study, this result suggests that further studies using extended sampling (infraspecific phylogenetic inferences or population genetics studies) are needed to clarify species boundaries (Figure 1).

*Microgramma mortoniana*, for example, was described as a hybrid between *M. squamulosa* and *M. vacciniifolia* (de la Sota, 1973b) and appears nested in the former species. The polyphyly of *Microgramma tobagensis* suggests a complex history where hybridization could play a role. The samples of *M. tobagensis* included in this study are from Peru, the main part of the species range, and eastern Brazil, a disjunct population that could also indicate either an undergoing allopatric speciation event or reticulation involving the sympatric *M. reptans*. *Microgramma tobagensis* is morphologically very similar to *M. reptans* (Smith *et al*., 2018) and it was considered a synonym of *M. piloselloides* by Tryon & Stolze (1993). However, according to the circumscription of Almeida (2014), *M. tobagensis* and *M. piloselloides* are not sympatric, while *M. tobagensis* distribution largely overlaps with that of *M. reptans* (Smith *et al*., 2018; Almeida, 2020). A comprehensive sampling, data at the population level, and other tools are needed to clarify the species circumscriptions and the evolutionary processes playing a role in these lineages.

*Microgramma microsoroides* is one of the most distinctive species in the genus by having irregularly scattered sori (Figure 3). All other *Microgramma* species present two rows, one row on each side of the costa, a character that was used to recognize the genus before its recircumscription (Salino *et al*., 2008). The presence of the irregularly scattered sori could be related to a hybrid origin, perhaps even an intergeneric one – events that are not rare among ferns (e.g. Rothfels *et al*., 2015; Engels & Canestraro, 2017). Alternatively, the hypothesis raised by Salino *et al*. (2008) that the multiple, scattered sori in *M. microsoroides* could have originated by an evolutionary breakup of linear sori – a feature present in *M. persicariifolia* (Figure 3) – should be further investigated given its close relationship with *M. persicariifolia*, a species with distinctive linear sori. More data, including nuclear data, population sampling, and chromosome counting are needed to investigate both the origin of *M. microsoroides* and the relations between these two species.

### Dating and ecological and biogeographical implications

To the date, there are no known fossils of *Microgramma* or closely related genera. To estimate the dates, we chose the secondary calibration points recovered by Testo & Sundue (2016). These authors used more Polypodiaceae fossils to calibrate their analysis and recovered older ages in this family than other studies did (e.g. Schuettpelz & Pryer, 2009) and therefore propose a more conservative premise (Testo & Sundue, 2016). The main clades inside *Microgramma* all diverged in the Neogene, around 9 – 15 Ma, except for the Scaly clade, which is older (ca. 23 Ma) (Figure 2). Most species divergence appears to have occurred in the Miocene or at the beginning of the Pliocene (Figure 2).

The position of *Microgramma mauritiana* (Figure 1, 2) nested within neotropical species indicates a long-distance dispersal event from America to Africa followed by speciation, as already mooted by Moran & Smith (2001). The LDD is estimated to have occurred around 15 Ma ago, long after the breakup of Gondwana (Jokat *et al*., 2003; Will & Frimmel, 2018) and the final separation of the African and South American continents, around 90 Ma ago (Thomaz Filho *et al*., 2000) (Figures 1, 2). This pattern has been observed in other ferns such as Thelypteridaceae (Almeida *et al*., 2016), *Ctenitis* (C.Chr.) C.Chr. (Hennequin *et al*., 2017), *Lellingeria* A.R.Sm. & R.C.Moran (Labiak, Sundue, & Rouhan, 2010) and *Platycerium* Desv. (Kreier & Schneider, 2006), among others. Examples of this pattern can also be found among lycophytes, such as *Phlegmariurus* Holub (Bauret *et al*., 2018), and angiosperms (Renner, 2004; Christenhusz & Chase, 2013).

Long-distance dispersal is a key subject in the biogeographic studies of ferns and lycophytes (Tryon, 1986; Barrington, 1993). LDD appears to have a major impact in the current distribution of the species-level lineages, as has been evidenced from this and other recent phylogenetic hypotheses, while vicariance likely had a major role in building biogeographic patterns in deep evolutionary scale for these plants (e.g. Lehtonen *et al*., 2010; Chao *et al*., 2014; Sundue *et al*., 2014; Labiak *et al*., 2015; Almeida *et al*., 2016; Hennequin *et al*., 2017; Vicent, Gabriel y Galán, & Sessa, 2017; Bauret *et al*., 2018). Divergence times of extant fern species from South America and African floras are younger than the Gondwana split and the separation of Africa and South America, so that at the specific level, the current disjunctions should reflect LDD rather than vicariant events (Renner, 2004).

Our dating estimates the origins of ant-fern association in *Microgramma* to be no older than 4.76 Ma in the lineage including *M. bifrons* and *M. brunei*. The divergence of *M. megalophylla* is older (11.67 Ma) but in this species, not all populations present signs of antassociation (Almeida, 2018). The absence in *M. megalophylla* of a specialized structure like the tuber-like myrmecodomatia of *M. bifrons* and *M. brunei* could indicate a very recent association with ants. Our results suggest a more recent evolution of ant association and myrmecodomatia in *Microgramma* compared to that proposed for *Lecanopteris* (16.70–21.80 Ma - Testo & Sundue, 2016).

The absence of geographic patterns inside the clades might be related to ecological preferences in this lineage. Most species in *Microgramma* are widespread or have a broad extent of occurrence, with few species considered narrow endemics (Almeida, 2014). They occur preferentially in elevations below 2000 m.a.s.l., with richness and abundance decreasing in higher elevations (Almeida, 2014; Smith *et al*., 2018). They are mainly epiphytes without great habitat specificities, usually presenting a large capacity of dispersal and colonization of a diverse array of habitats. These ecological aspects might obscure the biogeographic history of the genus or might indicate that speciation occurs mainly allopatrically in *Microgramma*, with dispersal inside the neotropical realm playing an important role in the evolution of the lineages inside the genus.

## CONCLUSIONS

The extensive sampling of *Microgramma* presented here supports the monophyly of the genus as currently circumscribed. Our results indicate that a considerable part of the broad morphological variation observed in the genus is homoplastic, as well as ecological interactions such as ant-association. The young age we found for the African lineage of *Microgramma* (15 Ma) corroborates a long-distance dispersal event from South America to Africa and reinforces its important role in the floristic relationships between South America and Africa.

## Acknowledgments

This work was supported by FAPEMIG (Fundação de Amparo à Pesquisa do Estado de Minas Gerais) through grant CRA-APQ 01599-10, CNPq (Conselho Nacional de Desenvolvimento Científico e Tecnológico) through grants 478723/2010-5 and 563568/2010, and for productivity grants, all attributed to AS, and the ATM “Biodiversité actuelle et fossile” of the Muséum national d’Histoire naturelle (2012) attributed to SH. CNPq also provided research scholarships to TEA (grant numbers 555226/2010-7 and 202160/2011-4). We thank the Coordenac□ão de Aperfeic□oamento Pessoal de Nível Superior (CAPES) for financing the 1st Scientific Publication Workshop of Programa de Pós-graduação em Biodiversidade through the PROCAD-AM grant 88887.200472/2018-00. We would like to thank the Muséum national d’Histoire naturelle de Paris, France (MNHN), the Service de Systématique Moléculaire (SSM - MNHN) and Céline Bonillo for the support and help. We also thank O. Álvarez-Fuentes, C.N. Fraga, M. Gaudeul, A.L. Gasper, L.L. Giacomin, J. Hickey, M. Kessler, M. Lautert, J. Lombardi, F. Matos, G. Rouhan, and F.S. Souza, for kindly sharing dried silica leaves or information that helped in the collections; Heidelberg Botanical Garden, Berlin Botanical Garden and Munich Botanical Garden, for permission for collect from cultivated plants; L.L. Giacomin, F. Costa, I. Kaefer, and A.R. Field for their contributions to this manuscript.

## APPENDIX Collection information for voucher specimens of sequences generated in this study and GenBank accession numbers^2^

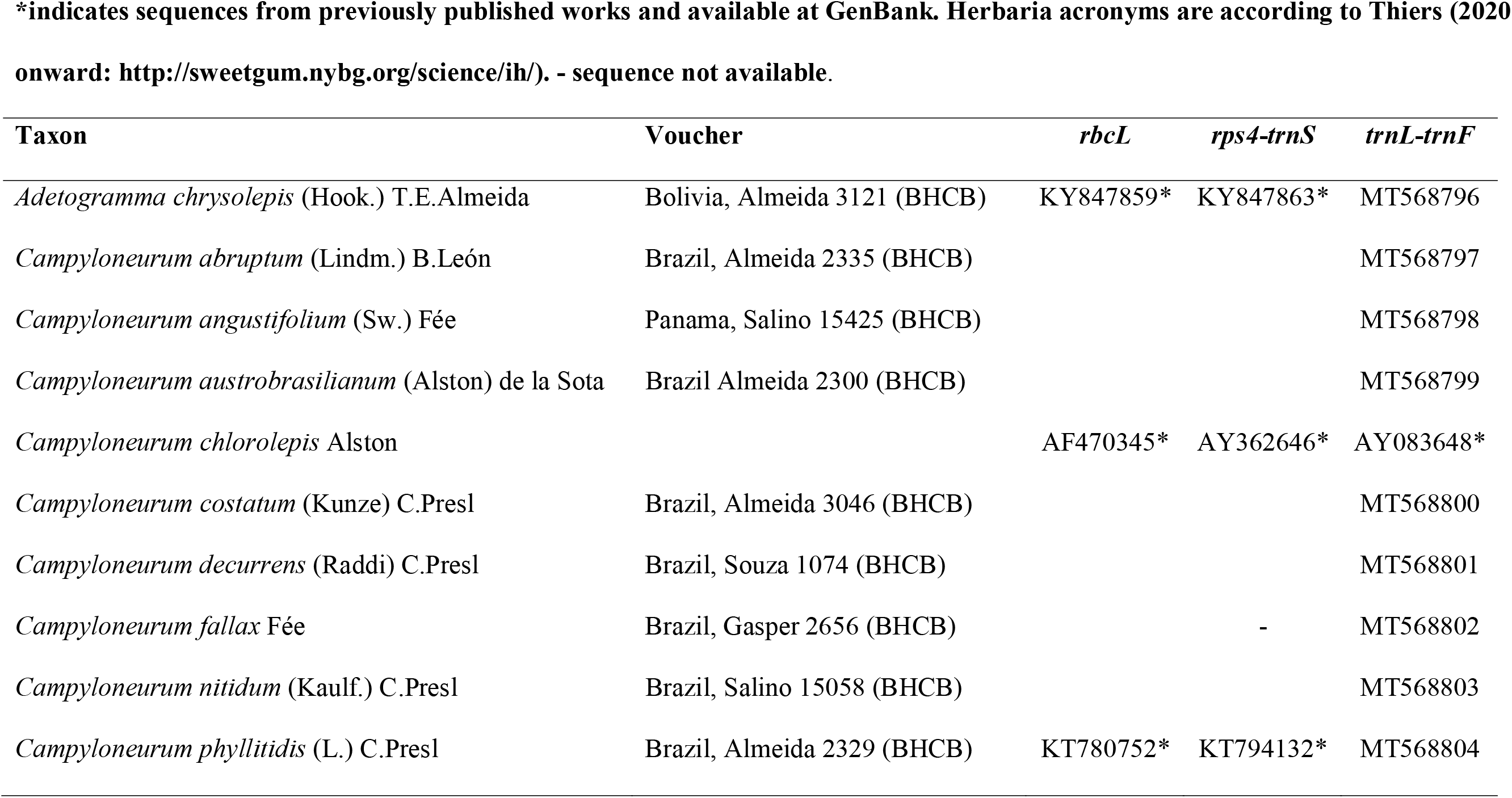
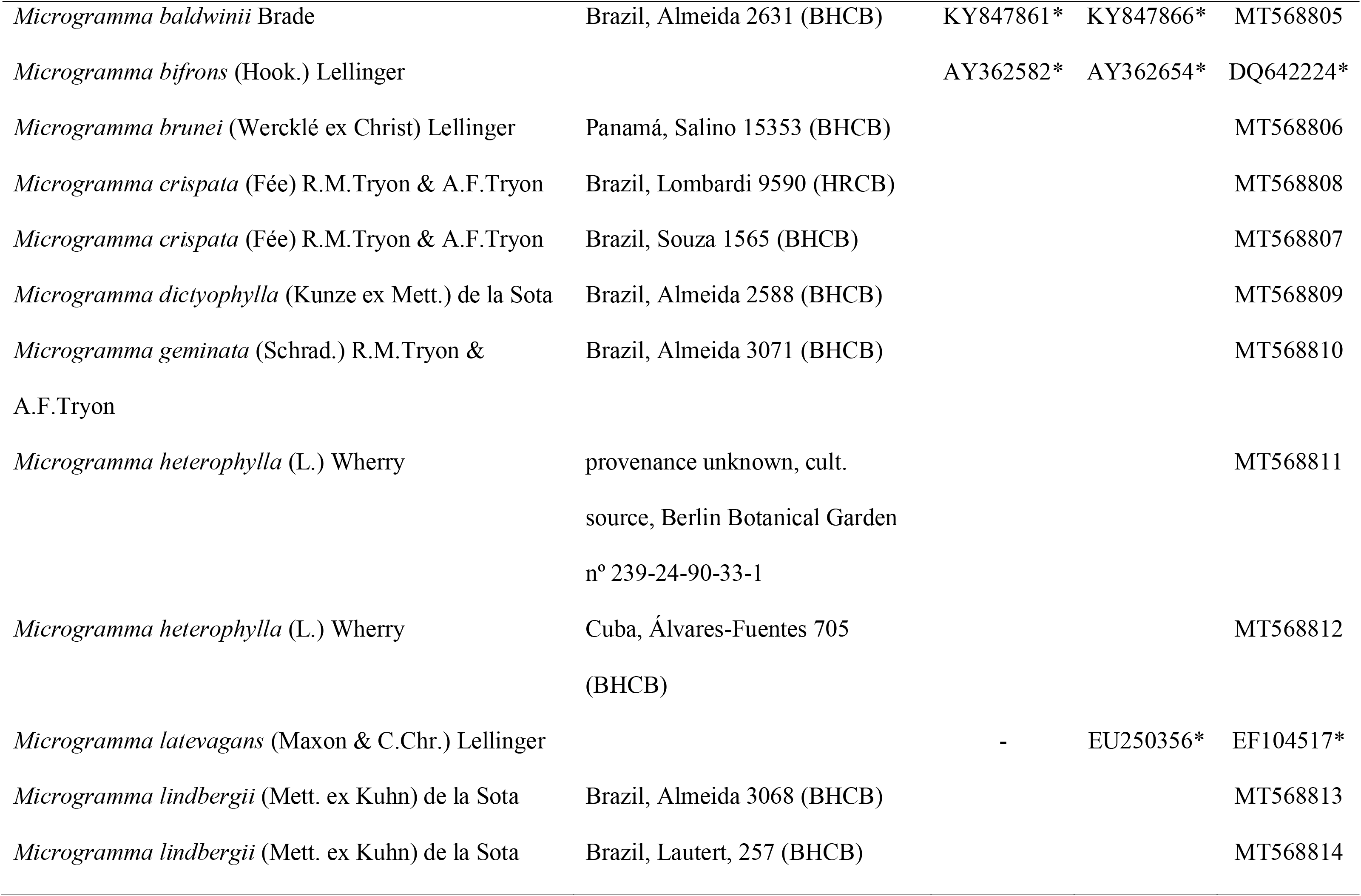
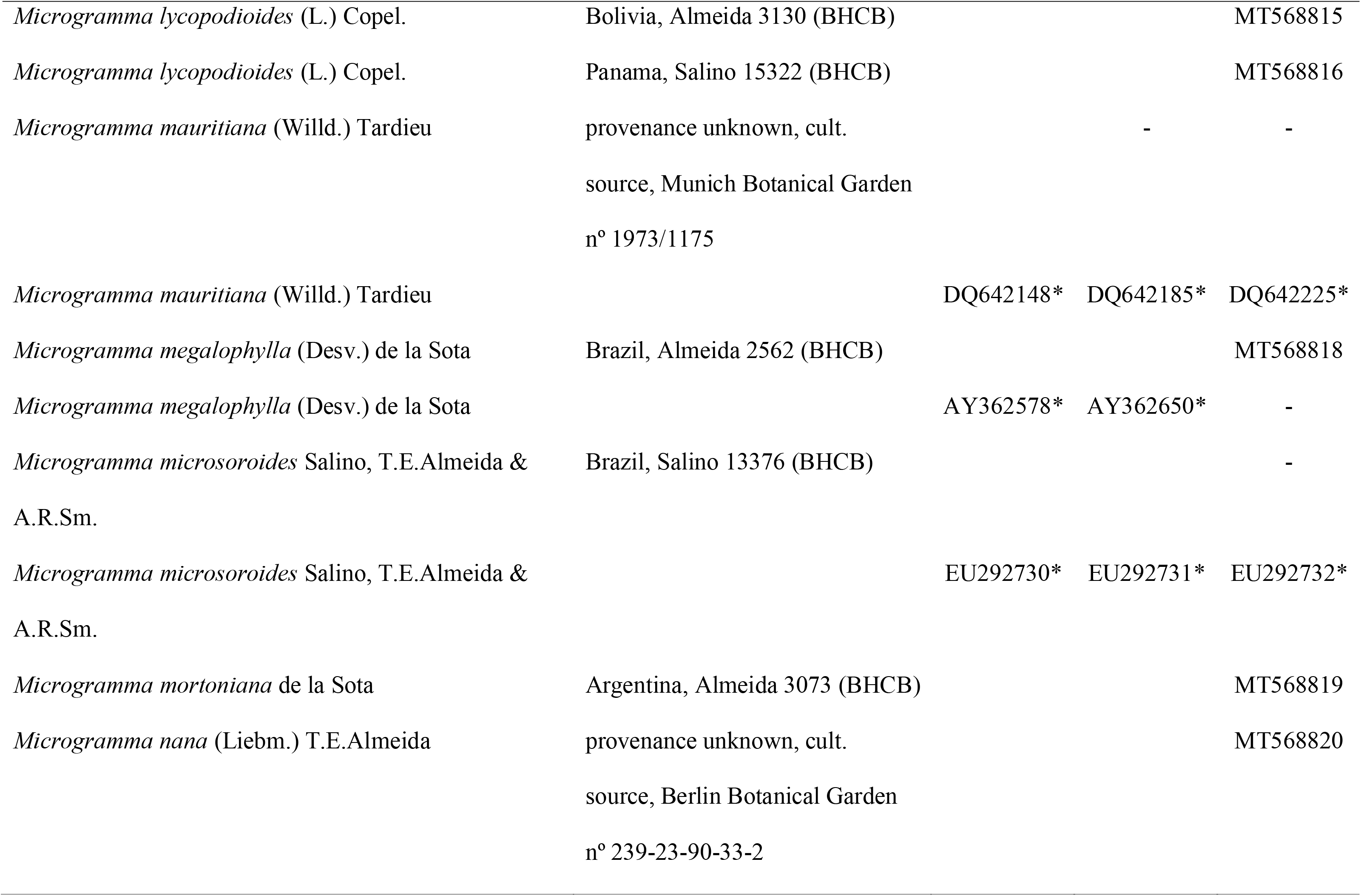
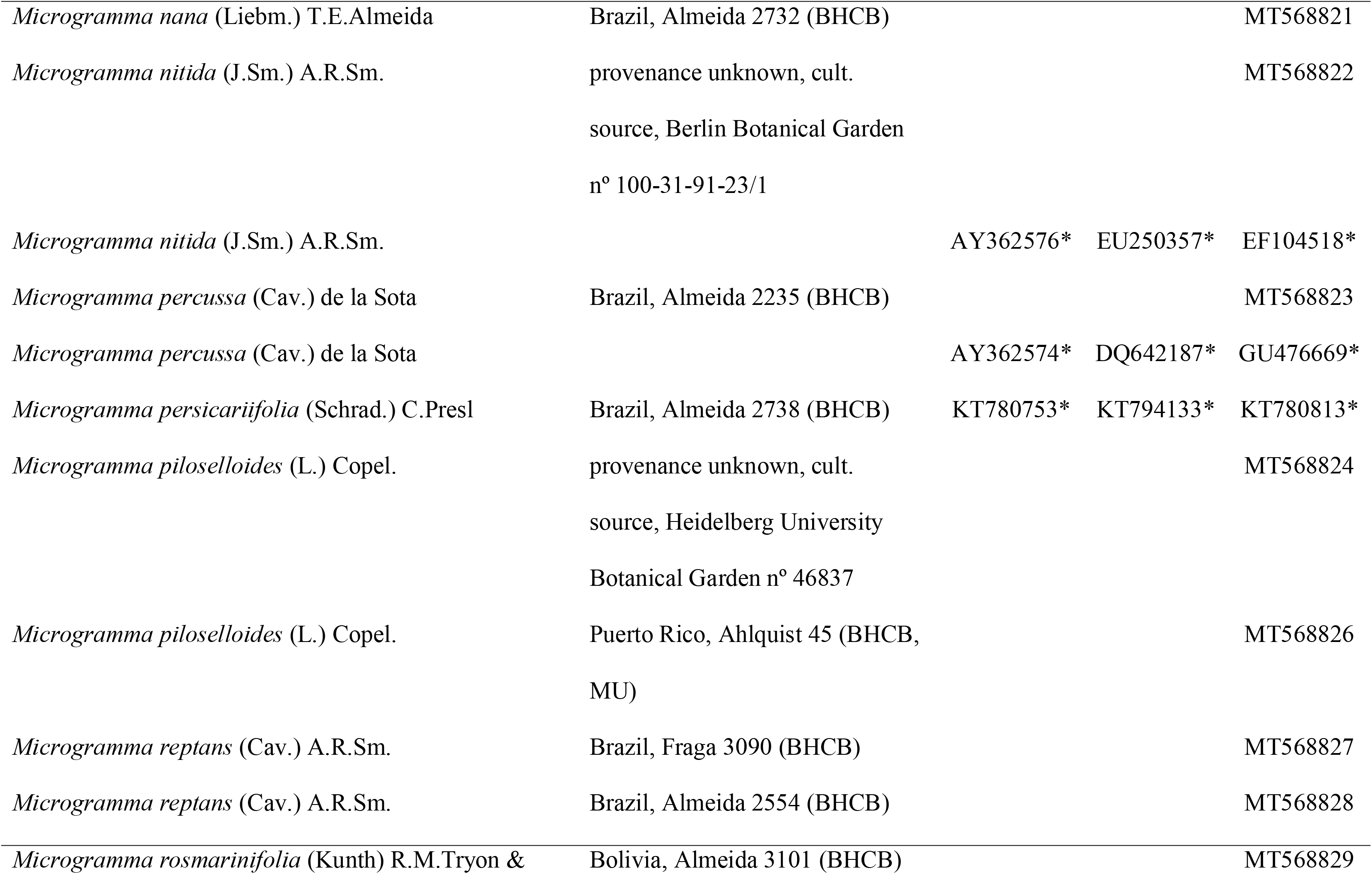
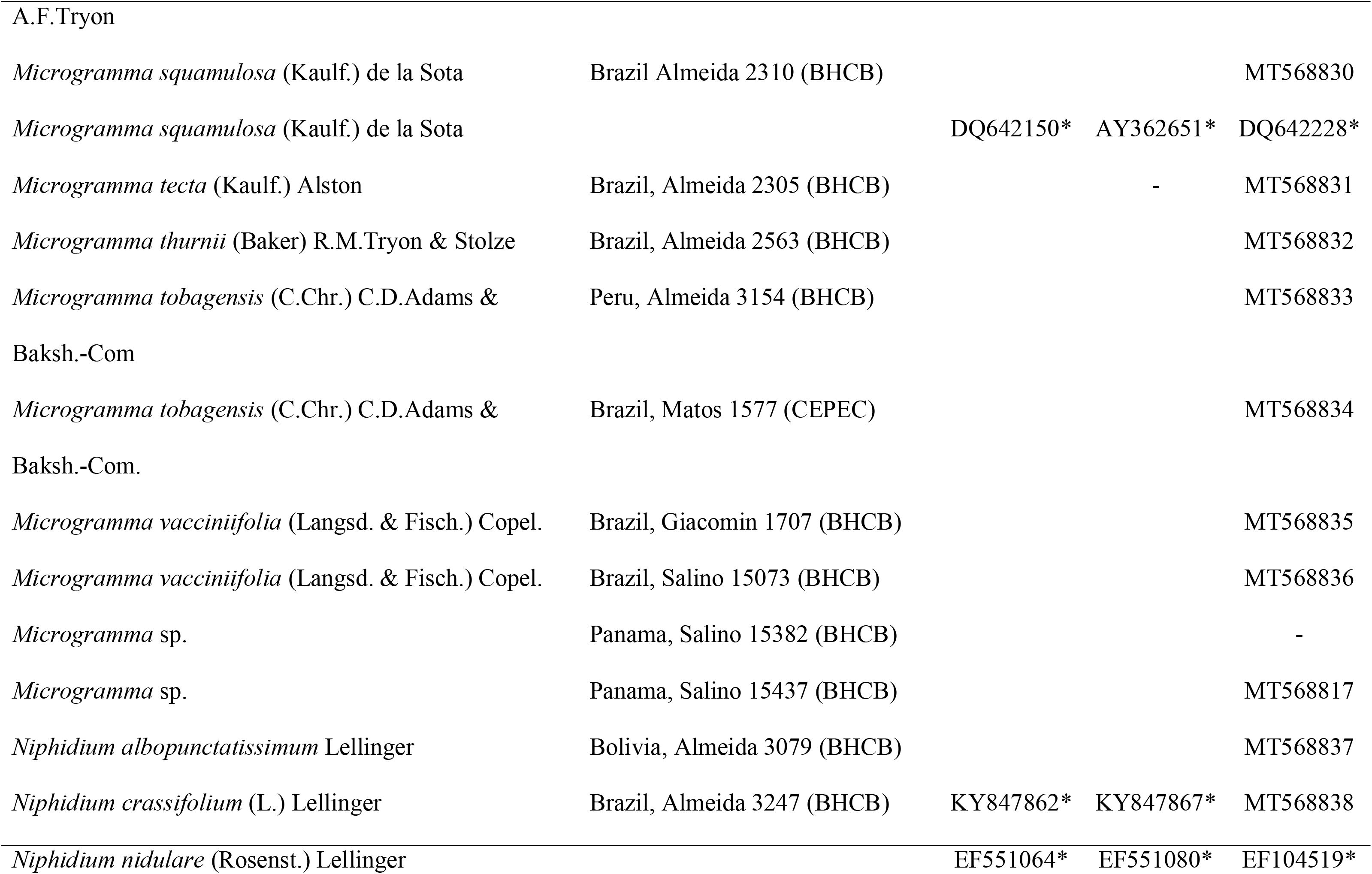
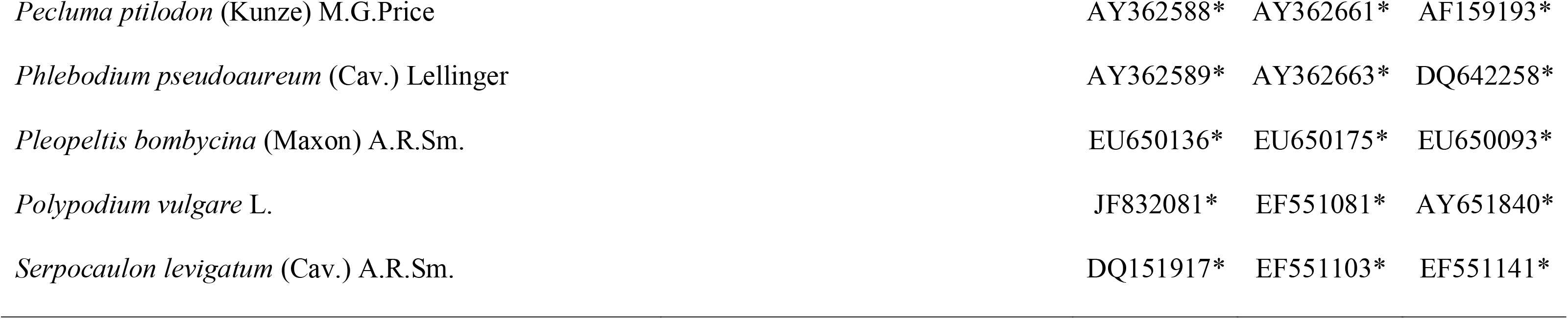

## Footnotes

2 We already submitted the sequences to GenBank and the accessions will be added as soon as they are sent.

## References

Almeida TE. 2014. Systematic studies in the genus *Microgramma* C.Presl (Polypodiaceae-Polypodiopsida).

Almeida TE, Hennequin S, Schneider H, Smith AR, Batista JAN, Ramalho AJ, Proite K & Salino A. 2016. Towards a phylogenetic generic classification of Thelypteridaceae: Additional sampling suggests alterations of neotropical taxa and further study of paleotropical genera. Molecular Phylogenetics and Evolution 94: 688–700.

Almeida TE, Salino A, Dubuisson JY & Hennequin S. 2017. *Adetogramma* (Polypodiaceae), a new monotypic fern genus segregated from *Polypodium*. PhytoKeys 78: 109–131.

Almeida TE. 2018. Ant-Fern association in *Microgramma megalophylla*. American Fern Journal 108: 62–64.

Almeida TE. 2020. Microgramma. Flora do Brasil 2020 under construction. Jardim Botânico do Rio de Janeiro.

Barido-Sottani J, Bošková V, Plessis L Du, Kühnert D, Magnus C, Mitov V, Müller NF, PecErska J, Rasmussen DA, Zhang C, Drummond AJ, Heath TA, Pybus OG, Vaughan TG & Stadler T. 2018. Taming the BEAST—A Community Teaching Material Resource for BEAST 2. Systematic Biology 67: 170–174.

Barrington DS. 1993. Ecological and historical factors in fern biogeography. Journal of Biogeography 20: 275–279.

Bauret L, Rouhan G, Hirai RY, Perrie L, Prado J, Salino A, Senterre B, Shepherd L, Sundue M, Selosse MA & Gaudeul M. 2017. Molecular data, based on an exhaustive species sampling of the fern genus *Rumohra* (Dryopteridaceae), reveal a biogeographical history mostly shaped by dispersal and several cryptic species in the widely distributed Rumohra adiantiformis. Botanical Journal of the Linnean Society 185: 463–481.

Bauret L, Field AR, Gaudeul M, Selosse MA & Rouhan G. 2018. First insights on the biogeographical history of *Phlegmariurus* (Lycopodiaceae), with a focus on Madagascar. Molecular Phylogenetics and Evolution 127: 488–501.

Bouckaert R, Heled J, Kühnert D, Vaughan T, Wu CH, Xie D, Suchard MA, Rambaut A & Drummond AJ. 2014. BEAST 2: A Software Platform for Bayesian Evolutionary Analysis. PLoS Computational Biology 10: 1–6.

Bouckaert R, Vaughan TG, Barido-Sottani J, Duchêne S, Fourment M, Gavryushkina A, Heled J, Jones G, Kühnert D, De Maio N, Matschiner M, Mendes FK, Müller NF, Ogilvie HA, du Plessis L, Popinga A, Rambaut A, Rasmussen D, Siveroni I, Suchard MA, Wu CH, Xie D, Zhang C, Stadler T & Drummond AJ. 2019. BEAST 2.5: An advanced software platform for Bayesian evolutionary analysis (M Pertea, Ed.). PLOS Computational Biology 15: e1006650.

Chao YS, Rouhan G, Amoroso VB & Chiou WL. 2014. Molecular phylogeny and biogeography of the fern genus *Pteris* (Pteridaceae). Annals of Botany 114: 109–124.

Chernomor O, von Haeseler A & Minh BQ. 2016. Terrace aware data structure for phylog-enomic inference from supermatrices. Systematic Biology 65: 997–1008.

Ching RG. 1940. On the natural classification of ferns. Sunyatsenia 5: 269–288.

Christenhusz MJM & Chase MW. 2013. Biogeographical patterns of plants in the Neotropics - dispersal rather than plate tectonics is most explanatory. Botanical Journal of the Linnean Society 171: 277–286.

Christensen C. 1938. Filicinae. In: Verdoon F, ed. Manual of Pteridology. The Hague: M. Nijhoff, 522–550.

Copeland EB. 1951. A New Genus of Ferns. American Fern Journal 41: 75.

Darriba D, Taboada GL, Doallo R & Posada D. 2015. jModelTest 2: more models, new heuristics and high-performance computing Europe PMC Funders Group. Nature Methods 9: 772.

Drummond AJ, Suchard MA, Xie D & Rambaut A. 2012. Bayesian Phylogenetics with BEAUti and the BEAST 1.7. Molecular Biology and Evolution 29: 1969–1973.

Duan YF, Hennequin S, Rouhan G, Bassuner B & Zhang LB. 2017. Taxonomic revision of the fern genus *Ctenitis* (Dryopteridaceae) from Africa and the Western Indian Ocean. Annals of the Missouri Botanical Garden 102: 3–86.

Engels ME & Canestraro BK. 2017. *×Cyclobotrya:* A new hybrid genus between Cyclodium and Polybotrya (Dryopteridaceae) from the Brazilian Amazon. Brittonia: 1–6.

Gasper AL de, Almeida TE, Dittrich VA de O, Smith AR & Salino A. 2016. Molecular phylogeny of the fern family Blechnaceae (Polypodiales) with a revised genus-level treatment. Cladistics 33: 1–18.

Gay H. 1993. Rhizome structure and evolution in the ant-associated epiphytic fern *Lecanopteris* Reinw. (Polypodiaceae). Botanical Journal of the Linnean Society 113: 135–160.

Gómez LD. 1974. Biology of the potato-fern *Solanopteris brunei*. Brenesia 28: 37–61.

Guindon S & Gascuel O. 2003. A Simple, Fast, and Accurate Algorithm to Estimate Large Phylogenies by Maximum Likelihood. Systematic Biology 52: 696–704.

Haufler CH, Grammer WA, Hennipman E, Ranker TA, Smith AR & Schneider H. 2003. of the ant-fern genus *Lecanopteris* (Polypodiaceae): testing phylogenetic hypotheses with DNA sequences. Systematic Botany 28: 217–227.

Haufler CH, Pryer KM, Schuettpelz E, Sessa EB, Farrar DR, Moran R, Schneller JJ, Watkins JE & Windham MD. 2016. Sex and the single gametophyte: Revising the homosporous vascular plant life cycle in light of contemporary research. BioScience 66: 928–937.

Haufler CH & Ranker TA. 1995. rbcL sequences provide phylogenetic insights among sister species of the fern genus Polypodium. American Fern Journal 85: 361.

Hennequin S, Rouhan G, Salino A, Duan YF, Lepeigneux MC, Guillou M, Ansell S, Almeida TE, Zhang LB & Schneider H. 2017. Global phylogeny and biogeography of the fern genus *Ctenitis* (Dryopteridaceae), with a focus on the Indian Ocean region. Molecular Phylogenetics and Evolution 112: 277–289.

Hennipman E. 1990. The significance of the SEM for character analysis of spores of Polypodiaceae. In: Claugher D, ed. Scanning Electron Microscopy in Taxonomy and Functional Morphology. Oxford: Clarendon Press, 23–44.

Hoang DT, Chernomor O, von Haeseler A, Minh BQ & Vinh LS. 2018. UFBoot2: Improving the Ultrafast Bootstrap Approximation. Molecular biology and evolution. Molecular Biology and Evolution 35: 518–522.

Huiet L, Li FW, Kao TT, Prado J, Smith AR, Schuettpelz E & Pryer KM. 2018. A worldwide phylogeny of *Adiantum* (Pteridaceae) reveals remarkable convergent evolution in leaf blade architecture. Taxon 67: 488–502.

Jokat W, Boebel T, König M & Meyer U. 2003. Timing and geometry of early Gondwana breakup. Journal of Geophysical Research: Solid Earth 108.

Jordano P. 2017. What is long-distance dispersal ? And a taxonomy of dispersal events. Journal of Ecology 105: 75–84.

Kalyaanamoorthy S, Minh BQ, Wong TKF, von Haeseler A & Jermiin LS. 2017. ModelFinder: fast model selection for accurate phylogenetic estimates. Nature Methods 14: 587–589.

Kearse M, Moir R, Wilson A, Stones-Havas S, Cheung M, Sturrock S, Buxton S, Cooper A, Markowitz S, Duran C, Thierer T, Ashton B, Mentjies P & Drummond A. 2012. Geneious Basic: an integrated and extendable desktop software platform for the organization and analysis of sequence dataNo Title. Bioinformatics 28: 1647–1649.

Kreier HP & Schneider H. 2006. Phylogeny and biogeography of the staghorn fern genus *Platycerium* (Polypodiaceae, Polypodiidae). American Journal of Botany 93: 217–225.

Kumar S, Stecher G & Tamura K. 2016. MEGA7: Molecular Evolutionary Genetics Analysis version 7.0 for bigger datasets. Molecular Biology and Evolution 33: 1870–1874.

de la Sota ER. 1973a. On the classification and phylogeny of the Polypodiaceae. 67 Suppl.: 229–244.

de la Sota ER. 1973b. A new species of *Microgramma* from Argentina. American Fern 63: 61–64.

Labiak PH, Sundue M, Rouhan G, Hanks JG, Mickel JT & Moran RC. 2014. Phylogeny and historical biogeography of the lastreopsid ferns (Dryopteridaceae). American Journal of Botany 101: 1207–1228.

Labiak PH, Mickel JT & Hanks JG. 2015. Molecular phylogeny and character evolution of Anemiaceae (Schizaeales). Taxon 64: 1141–1158.

Labiak PH & Moran RC. 2018. Phylogeny of *Campyloneurum* (polypodiaceae). International Journal of Plant Sciences 179: 36–49.

Labiak PH, Sundue M & Rouhan G. 2010. Molecular phylogeny, character evolution, and biogeography of the grammitid fern genus *Lellingeria* (Polypodiaceae). American journal of botany 97: 1354–64.

Lehtonen S, Tuomisto H, Rouhan G & Christenhusz MJM. 2010. Phylogenetics and classification of the pantropical fern family Lindsaeaceae. Botanical Journal of the Linnean Society 163: 305–359.

Lellinger DB. 1977. Nomenclatural notes on some ferns of Costa Rica, Panama, and Colombia. American Fern Journal 67: 58–60.

León B & Jørgensen PM. 1999. Polypodiaceae. In: Jørgensen PM, León-Yánez S, eds. Catalogue of the vascular plants of Ecuador. Saint Louis: Missouri Botanical Garden Press, 154–168.

Link-Pérez MA, Watson LE & Hickey RJ. 2011. Redefinition of *Adiantopsis* Fée (Pteridaceae): Systematics, diversification, and biogeography. Taxon 60: 1255–1268.

Mehltreter K & Sharpe JM. 2013. Causes and consequences of the variability of leaf lifespan of ferns. Fern Gazette 19: 193–202.

Miller MA, Pfeiffer W & Schwartz T. 2010. Creating the CIPRES Science Gateway for inference of large phylogenetic trees. In: 2010 Gateway Computing Environments Workshop (GCE). IEEE, 1–8.

Minh BQ, Nguyen MAT & Von Haeseler A. 2013. Ultrafast approximation for phylogenetic bootstrap. Molecular Biology and Evolution 30: 1188–1195.

Moran RC & Riba R. 1995. Flora mesoamericana (G Davidse, M Sousa S., and S Knapp, Eds.). Ciudad del México: Universidad Autonoma de Mexico.

Moran RC & Smith AR. 2001. Phytogeographic Relationships between Neotropical and African-Madagascan Pteridophytes. Brittonia 53: 304–351.

Nadot S, Bittar G, Carter L, Lacroix R & Lejeune B. 1995. A phylogenetic analysis of monocotyledons based on the chloroplast gene rps4, using parsimony and a new numerical phenetics method. Molecular Phylogenetics and Evolution 4: 257–282.

Nguyen LT, Schmidt HA, von Haeseler A & Minh BQ. 2015. IQ-TREE: A fast and effective stochastic algorithm for estimating maximum likelihood phylogenies. Molecular Biology and Evolution 32: 268–274.

Paradis E & Schliep K. 2019. ape 5.0: an environment for modern phylogenetics and evolutionary analyses in R (R Schwartz, Ed.). Bioinformatics 35: 526–528.

Pichi Sermolli REG. 1977. Tentamen Pteridophytorum genera in taxonomicum ordinem redigendi. Webbia 31: 313–512.

Pinson JB, Chambers SM, Nitta JH, Kuo LYY & Sessa EB. 2017. The separation of generations: Biology and biogeography of long-lived sporophyteless fern gametophytes. International Journal of Plant Sciences 178: 1–18.

Prado J, Schuettpelz E, Hirai RY & Smith AR. 2013. *Pellaea flavescens*, a brazilian endemic, is a synonym of Old World *Pellaea viridis*. American Fern Journal 103: 21–26.

R Core Team. 2019. R: A Language and Environment for Statistical Computing.

Rambaut A, Drummond AJ, Xie D, Baele G & Suchard MA. 2018. Posterior Summarization in Bayesian Phylogenetics Using Tracer 1.7. Systematic Biology 67: 901–904.

Renner S. 2004. Plant Dispersal across the Tropical Atlantic by Wind and Sea Currents. International Journal of Plant Sciences 165: S23–S33.

Ronquist F, Teslenko M, Van Der Mark P, Ayres DL, Darling A, Höhna S, Larget B, Liu L, Suchard MA & Huelsenbeck JP. 2012. MrBayes 3.2: Efficient bayesian phylogenetic inference and model choice across a large model space. Systematic Biology 61: 539–542.

Rothfels CJ, Johnson AK, Hovenkamp PH, Swofford DL, Roskam HC, Fraser-Jenkins CR, Windham MD & Pryer KM. 2015. Natural hybridization between genera that diverged from each other approximately 60 million years ago. American Naturalist 185: 433–442.

Rouhan G, Dubuisson JY, Rakotondrainibe F, Motley TJ, Mickel JT, Labat JN & Moran RC. 2004. Molecular phylogeny of the fern genus *Elaphoglossum* (Elaphoglossaceae) based on chloroplast non-coding DNA sequences: contributions of species from the Indian Ocean area. Molecular phylogenetics and evolution 33: 745–63.

Roux JP. 2009. Synopsis of the Lycopodiophyta and Pteridophyta of Africa, Madagascar and neighbouring islands. Pretoria: South African National Biodiversity Institute.

Salino A, Almeida TE, Smith AR, Gómez AN, Kreier HP & Schneider H. 2008. A New Species of *Microgramma* (Polypodiaceae) from Brazil and Recircumscription of the Genus Based on Phylogenetic Evidence. Systematic Botany 33: 630–635.

Schneider H, Smith AR, Cranfill R, Hildebrand TJ, Haufler CH & Ranker TA. 2004. Unraveling the phylogeny of polygrammoid ferns (Polypodiaceae and Grammitidaceae): Exploring aspects of the diversification of epiphytic plants. Molecular Phylogenetics and Evolution 31: 1041–1063.

Schuettpelz E, Chen CW, Kessler M, Pinson JB, Johnson G, Davila A, Cochran AT, Huiet L & Pryer KM. 2016. A revised generic classification of vittarioid ferns (Pteridaceae) based on molecular, micromorphological, and geographic data. Taxon 65: 708–722.

Schuettpelz E & Pryer KM. 2009. Evidence for a Cenozoic radiation of ferns in an angiosperm-dominated canopy. Proceedings of the National Academy of Sciences 106: 11200–11205.

Schwarz G. 1978. Estimating the dimension of a model. Annals of Statistics 6: 461–464.

Sheffield E. 2008. Alternation of generations. In: Ranker TA, Haufler CH, eds. Biology and Evolution of Ferns and Lycophytes. New York: Cambridge University Press, 49–74.

Smith AR, Kessler M, León B, Almeida TE, Jiménez-Pérez I & Lehnert M. 2018. Prodromus of a fern flora for Bolivia. XL. Polypodiaceae. Phytotaxa 354: 1.

Smith AR & Cranfill RB. 2002. Intrafamilial Relationships of the Thelypteroid Ferns (Thelypteridaceae). American Fern Journal 92: 131.

Smith AR & Tejero-Díez JD. 2014. *Pleopeltis* (Polypodiaceae), a redifinition of the genus and nomenclatural novelties. Botanical Sciences 92: 43–58.

Sundue MA, Parris BS, Ranker TA, Smith AR, Fujimoto EL, Zamora-Crosby D, Morden CW, Chiou WL, Chen CW, Rouhan G, Hirai RY & Prado J. 2014. Global phylogeny and biogeography of grammitid ferns (Polypodiaceae). Molecular Phylogenetics and Evolution 81:195–206.

Taberlet P, Gielly L, Pautou G & Bouvet J. 1991. Universal primers for amplification of three non-coding regions of chloroplast DNA. Plant molecular biology 17: 1105–9.

Testo WL, Field AR, Sessa EB & Sundue M. 2019. Phylogenetic and morphological analyses support the resurrection of *Dendroconche* and the recognition of two new genera in Polypodiaceae subfamily Microsoroideae. Systematic Botany 44: 737–752.

Testo W & Sundue M. 2014. Primary Hemiepiphytism in *Colysis ampla* (Polypodiaceae) Provides New Insight into the Evolution of Growth Habit in Ferns. International Journal of Plant Sciences 175: 526–536.

Testo W & Sundue M. 2016. A 4000-species dataset provides new insight into the evolution of ferns. Molecular Phylogenetics and Evolution 105: 200–211.

Thomaz Filho A, Mizusaki AMP, Milani EJ & Cesero P de. 2000. Rifting and magmatism associated with the South America and Africa break up. Revista Brasileira de Geociências 30: 017–019.

Trewick S, Morgan-Richards M, Russell SJ, Henderson S, Rumsey FJ, Pintér I, Barrett JA, Gibby M & Vogel JC. 2002. Polyploidy, phylogeography and Pleistocene refugia of the rockfern *Asplenium ceterach:* evidence from chloroplast DNA. Molecular ecology 11: 2003–12.

Trifinopoulos J, Nguyen LT, von Haeseler A & Minh BQ. 2016. W-IQ-TREE: a fast online phylogenetic tool for maximum likelihood analysis. Nucleic acids research 44: W232–W235.

Tryon R. 1986. The Biogeography of Species, with Special Reference to Ferns. The Botanical Review 52: 117–156.

Tryon AF & Lugardon B. 1991. Spores of the Pteridophyta. New York: Springer-Verlag.

Tryon RM & Stolze RG. 1993. Pteridophyta of Peru. Part V. 18. Aspleniaceae - 21. Polypodiaceae. Botany, new series 32: 1–190.

Vicent M, Gabriel y Galán JM & Sessa EB. 2017. Phylogenetics and historical biogeography of *Lomaridium* (Blechnaceae: Polypodiopsida). Taxon 66: 1304–1316.

Wagner WH & Wagner FS. 1977. Fertile-sterile leaf dimorphy in Ferns. The Gardens’ Bulletin Singapore 30: 251–267.

Watkins JE, Churchill AC & Michele Holbrook N. 2016. A site for sori: Ecophysiology of fertile-sterile leaf dimorphy in ferns. American Journal of Botany 103: 845–855.

Will TM & Frimmel HE. 2018. Where does a continent prefer to break up? Some lessons from the South Atlantic margins. Gondwana Research 53: 9–19.

